# CLASS-II KNOX genes coordinate spatial and temporal patterns of the tomato ripening

**DOI:** 10.1101/2021.11.19.469310

**Authors:** Alexandra Keren-Keiserman, Amit Shtern, Daniel Chalupowicz, Chihiro Furumizu, John Paul Alvarez, Ziva Amsalem, Tzahi Arazi, Sharon Tuvia-Alkalai, Idan Efroni, Elazar Fallik, Alexander Goldshmidt

## Abstract

Ripening is a complex developmental change of a mature organ, the fruit. In plants like a tomato, it involves softening, pigmentation, and biosynthesis of metabolites beneficial for the human diet. Examination of the transcriptional changes towards ripening suggests that redundant uncharacterized factors may be involved in the coordination of the ripening switch. Previous studies have demonstrated that Arabidopsis *CLASS-II KNOX* genes play a significant role in controlling the maturation of siliques and their transition to senescence. Here we examined the combined role of all four tomato *CLASS-II KNOX* genes in the maturation and ripening of fleshy fruits using an artificial microRNA targeting them simultaneously. As expected, the knockdown plants (*35S::amiR-TKN-CL-II*) exhibited leaves with increased complexity, reminiscent of the leaf phenotype of plants overexpressing *CLASS-I KNOX*, which antagonize *CLASS-II KNOX* gene functions. The fruits of *35S::amiR-TKN-CL-II* plants were notably smaller than the control. While their internal gel/placenta tissue softened and accumulated the typical pigmentation, the pericarp color break took place ten days later than control, and eventually, it turned yellow instead of red.

Additionally, the pericarp of *35S::amiR-TKN-CL-II* fruits remained significantly firmer than control even after three weeks of shelf storage. Strikingly, the *35S::amiR-TKN-CL-II* fruits showed early ethylene release and respiration peak, but these were correlated only with liquefaction and pigmentation of the internal tissues. Our findings suggest that *CLASS-II KNOX* genes are required to coordinate the spatial and temporal patterns of tomato fruit ripening.

**One sentence summary:** Tomato *CLASS-II KNOX* genes play antagonistic roles in the regulation of ripening at the internal fruit domains and pericarp.

## Introduction

Fleshy fruits are an essential component of the human diet. The unique feature of the fleshy fruits is that upon the termination of fruit set and growth, they begin to ripen (Seymour et al., 2013; Karlova et al., 2014). Ripening is defined by the fruit ability to undergo complex developmental changes through which it is softening, changing its metabolic content, color, nutritional levels, and aroma (Klee and Giovannoni 2011; Seymour et al., 2013; Li et al., 2018; Wang et al., 2018; Fenn and Giovannoni 2020; Wang et al., 2020). Examination of transcriptional and metabolic changes associated with the transition to ripening suggests that redundant unidentified genes and complex hormonal networks coordinate this process (Shinozaki et al., 2018; Fenn and Giovannoni 2020; Wang et al., 2020). It was shown that ripening initiation and progression are regulated differently in the two primary physiological fleshy fruit groups: climacteric and non-climacteric. In climacteric fruits, the process is associated with a sharp increase of respiration and synthesis of the phytohormone ethylene (climacteric peak), whereas in non-climacteric fruits, it is not linked to these two activities (Klee and Giovannoni 2011; Seymour et al., 2013; Li et al., 2018; Fenn and Giovannoni 2020).

Ethylene was also shown to promote leaf and dry fruit senescence, suggesting that similar mechanisms may coordinate the initiation of senescence and ripening (Koyama 2014; Khan et al., 2014; Bakshi et al., 2015; Iqbal et al., 2017). Comparative transcriptome analysis of *Arabidopsis thaliana* senescing dry siliques (fruit) and tomato (*Solanum lycopersicum*) ripening climacteric fruits have also indicated the existence of parallel maturation programs in both species (Gomez et al., 2014). However, the molecular players coordinating fruit maturation programs in dry and fleshy fruits have not yet been identified (Wang et al., 2020).

Previous studies have demonstrated that *Arabidopsis CLASS-II KNOX* genes redundantly regulate lateral organ development and maturation, including siliques growth and transition to senescence (Furumizu et al., 2015). The current study used artificial microRNA targeting all members of the tomato *CLASS-II KNOX* gene-clade to examine its functions in the development and ripening of fleshy fruits. Our results demonstrate that the ubiquitous knockdown of the *SlCLASS-II KNOX* genes increases leaf complexity, a phenotype reminiscent of leaves overexpressing their *CLASS-I KNOX* antagonists (Hareven et al., 1996; Janssen et al., 1998). The fruits of the *SlCLASS-II KNOX* knockdown plants were notably smaller than the control. Their internal domains (locular and placental tissues) exhibited evidence of ripening; however, their pericarp tissues did not accumulate red color or soften. Examination of the *SlCLASS-II KNOX* knockdown fruit ethylene and carbon dioxide emission levels indicated that the fruits reached the climacteric peak earlier than the controls and maintained ethylene levels similar to controls at the later stages. These findings demonstrate that *SlCLASS-II KNOX* genes regulate fruit maturation programs in both dry and fleshy fruits and coordinate spatial and temporal ripening patterns in tomato fruits.

## Results

### Tomato *CLASS-II KNOX* genes mediate the development of lateral organs

To explore the roles of the *CLASS-II KNOX* genes in tomato, we first identified the orthologs of the *Arabidopsis thaliana CLASS-II KNOX* genes (*TKN-CL-II*) (Furumizu et al., 2015). Phylogenetic analysis divided the CLASS-II KNOX proteins into a KNAT3/4/5*-*like clade and a KNAT7-like clade. While the KNAT7-like clade has a single gene in both *Arabidopsis* and tomato, the KNAT3/4/5 clade consists of three genes in both species, but these likely arise from independent duplications. For consistency with *Arabidopsis* nomenclature, we named the tomato KNAT7-like gene *SlKNAT7* and the three tomato *SlKNAT3*-like genes *SlKNAT3, SlKNAT4*, and *SlKNAT5* (Fig. 1A). We then used Tomato Expression Atlas (TEA) (http://tea.solgenomics.net; Shinozaki et al., 2018) to monitor fruit expression patterns of these genes. We found that *SlKNAT3, SlKNAT5*, and *SlKNAT7* genes are broadly expressed throughout fruit tissues (Fig. S1). In contrast, the *SlKNAT4* gene has low expression levels during fruit development that increased slightly at the post-breaker stages in the placenta, columella, and locular tissues (Fig S1).

**Figure 1:**
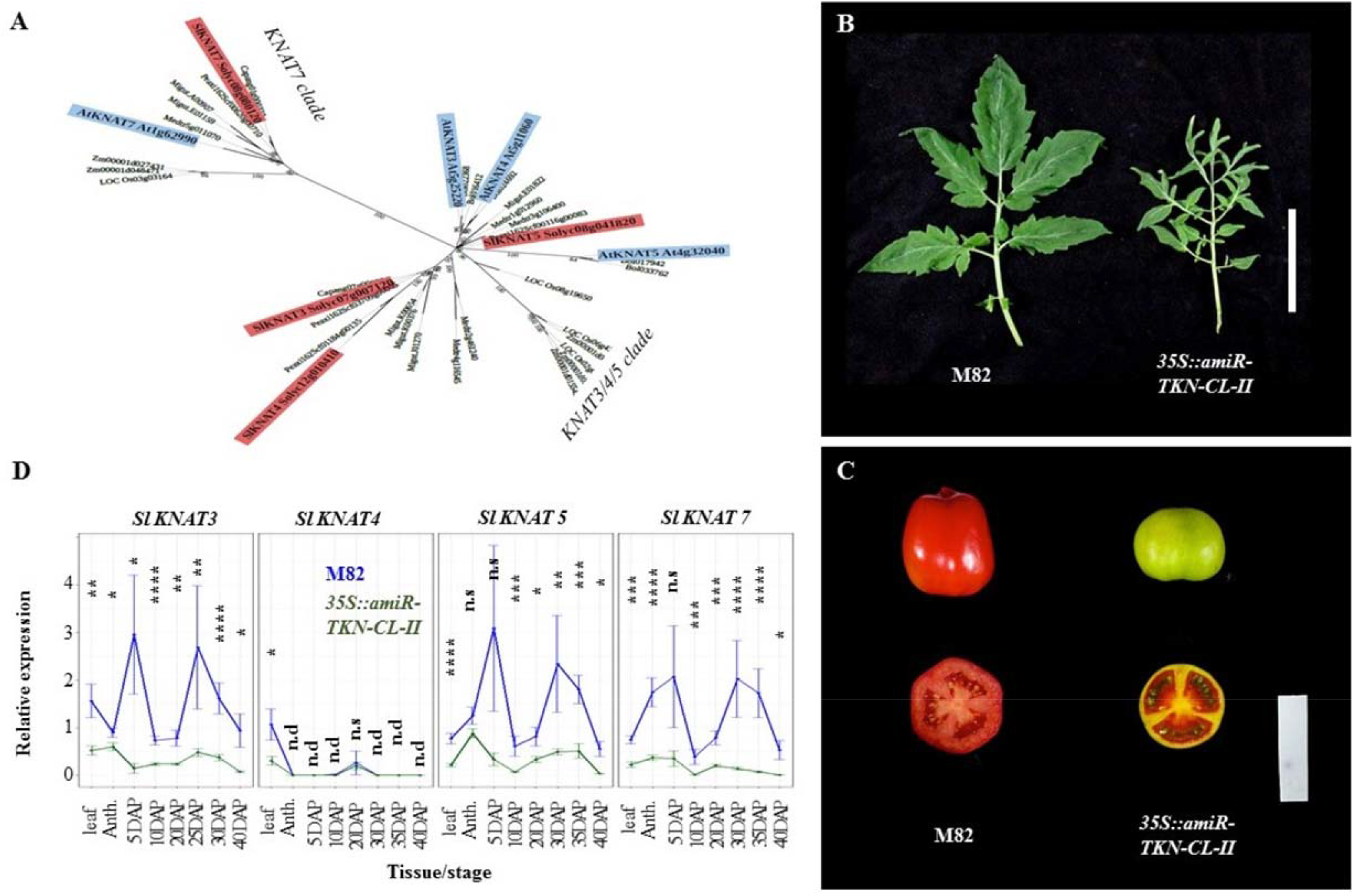
The tomato *CLASS-II KNOX* genes regulate fruit ripening and development. **A:** Phylogenetic tree showing homology, reflected by branch length and by numbered bootstrap values, among the *Arabidopsis thaliana* (blue) and *Solanum lycopersicum* (red) CLASS II KNOX proteins. The two distinct protein clades found in the gene family are named after *Arabidopsis* gene names. **B**: Forth leaves of M82 and *CLASS-II KNOX* knockdown line. The mutant leaves are more dissected, and leaflets are narrow. **C:** Whole and cut fruits images taken at 50 DAP when M82 fruits reach the Ripen Red (R. R.) stage. Scale bar in B-C is 6cm. **D:** Relative expression levels (Y-axis) of tomato *CLASS-II KNOX* genes in the fourth leaf, carpels at anthesis, 5 Days After Pollination (DAP) fruits,10, 20,25,30, and 40 DAP pericarps (X-axis) of M82 (blue) and *35S::amiR-TKN-CL-II* transgenic line (dark-green); dots indicate the mean (Relative expression) values; the error bars indicate standard errors. Student’s t-test H_0_ probabilities for each developmental time point are indicated as (**n.s**) (p-value)> 0.05; (**\***) (p-value) ≤ 0.05; (**\*\***) (p-value) ≤ 0.01; (**\*\*\***) (p-value) ≤ 0.001; (**\*\*\*\***) (p-value) ≤ 0.0001; **n.d** – indicates non detectable levels.

To test the function of the *CLASS-II KNOX* genes, we designed two artificial microRNAs (amiRs) using the *Arabidopsis* miR159a as a backbone. One amiR specifically targeted the tomato *SlKNAT3, SlKNAT4, SlKNAT5*, and the other amiR was specific to *SlKNAT7* (Alvarez et al., 2006; Fig. S2A). Both amiRs were cloned in tandem (Fig. S2B) either under the control of the 35S promoter or the bacterial OPERATOR (OP) promoter (Moore et al., 1998) and then transformed into tomato plants. Upon recovery of the *35S::amiR-TKN-CL-II* T0 transgenic progeny, increased leaf complexity was observed in multiple independent plants (Fig. 1B). These vegetative phenotypes appeared similar to those of the *A. thaliana class-II knox* mutants, transgenic plants expressing similarly designed amiR targeting these *AtKNOX-CL-II* genes (Furumizu et al., 2015) as well as to tomato plants ectopically expressing *CLASS-I KNOX* genes (Hareven et al., 1996; Janssen et al., 1998). A single weak T0 event was chosen as a working line due to the fact that stronger T0 plants produced very little pollen, such that no progeny could be recovered. Comparable to the observations reported for the fruit (siliques) of the *Arabidopsis class-II knox* mutants (Furumizu et al., 2015), the size of the *35S::amiR-TKN-CL-II* fruits was noticeably smaller than the control; in addition, the transgenic line fruits had an oblate shape versus the prolate shape of M82 fruits (Fig. 1C). By 50 Days After Pollination (DAP), M82 fruits reached the Ripe Red (RR) stage; at that age, the locular tissues of the *35S::amiR-TKN-CL-II* fruits were also fully ripened, whereas the fruit pericarp and septum tissues turned yellow instead of red (Fig. 1C).

The weak *35S::amiR-TKN-CL-II* event progeny was verified for a single insert at T1 generation. Like the M82 control, these plants flowered after producing eight to nine leaves (Fig. S3A) however, they developed abnormalities similar to those observed at independent T0 plants. q-RT-PCR analysis indicated that these phenotypes were associated with reduced *SlCLASS-II KNOX* transcription levels. Eight different tissues, including the fourth - leaf, carpels at anthesis, immature fruits at 5-DAP, and 10, 20, 25, 30, 40 (DAP) developing pericarps, were collected from the transgenic and M82 (control) plants. We detected significant downregulation of all four *SlCLASS-II KNOX* family members in the fourth leaf of the transgenic plants (Fig. 1D). Only *SlKNAT3* and *SlKNAT7* transcripts were significantly downregulated at the Anthesis-stage carpels, and just *SlKNAT3* transcripts were significantly reduced at the 5DAP transgenic fruits. Further on at the 10, 20, 25,30, and 40 DAP - transgenic pericarps, the transcription levels of the *SlKNAT3, SlKNAT7*, and *SlKNAT5* genes were significantly reduced. Due to its barely detectable transcription levels (Fig. S1), we could not reliably detect or assess the relative differences in the *SlKNAT4* expression in immature fruits (Fig. 1C). Taken together, our results indicate that the pleiotropic phenotypes observed in the *35S::amiR-TKN-CL-II* line were caused by reduced *SlCLASS-II KNOX* transcript levels (Fig. 1B-D).

### The *SlCLASS-II KNOX* genes regulate fruit size and ripening patterns

To characterize in detail the role of the *TKN-CL-II* genes in tomato fruit development, we compared M82 and *35S::amiR-TKN-CL-II* fruits along with their development and ripening (Fig. 2A). Significant differences were not observed until the 10DAP stage. By 20 DAP, the *SlCLASS-II KNOX* knockdown fruits were notably lighter and less elongated than the control (Fig. 2A-B). The weight of the *35S::amiR-TKN-CL-II* fruits continued to increase until 35DAP but always remained lower than control fruits, which continued to grow until 40DAP (Fig. 2A-B). Visual assessment of the fruit maturation patterns has detected that at 40DAP, M82 fruits reached the “Breaker” (BR) stage, at which initial patches of color change were visible at the fruit blossom-end, placenta, and very mildly in the locular tissues. At the same time (40DAP), no color change was observed in the exterior of the *35S::amiR-TKN-CL-II* fruits, whereas the internal (locular and placental) tissues showed noticeably advanced coloration than the control (Fig. 2A). Following the BR stage, M82 fruits progressed through the ripening stages (defined by the fruit color change), reaching the “Light Red” (LR) stage at about 46 DAP and “ Red Ripe” (RR) by the 50DAP. The locular and placental tissues of the *35S::amiR-TKN-CL-II* fruits progressed through ripening comparably to the M82 fruits, however, the pericarp and septum tissues of these fruits maintained the green color until 50DAP, after which they gradually turned yellow (Fig 2A).

**Figure 2:**
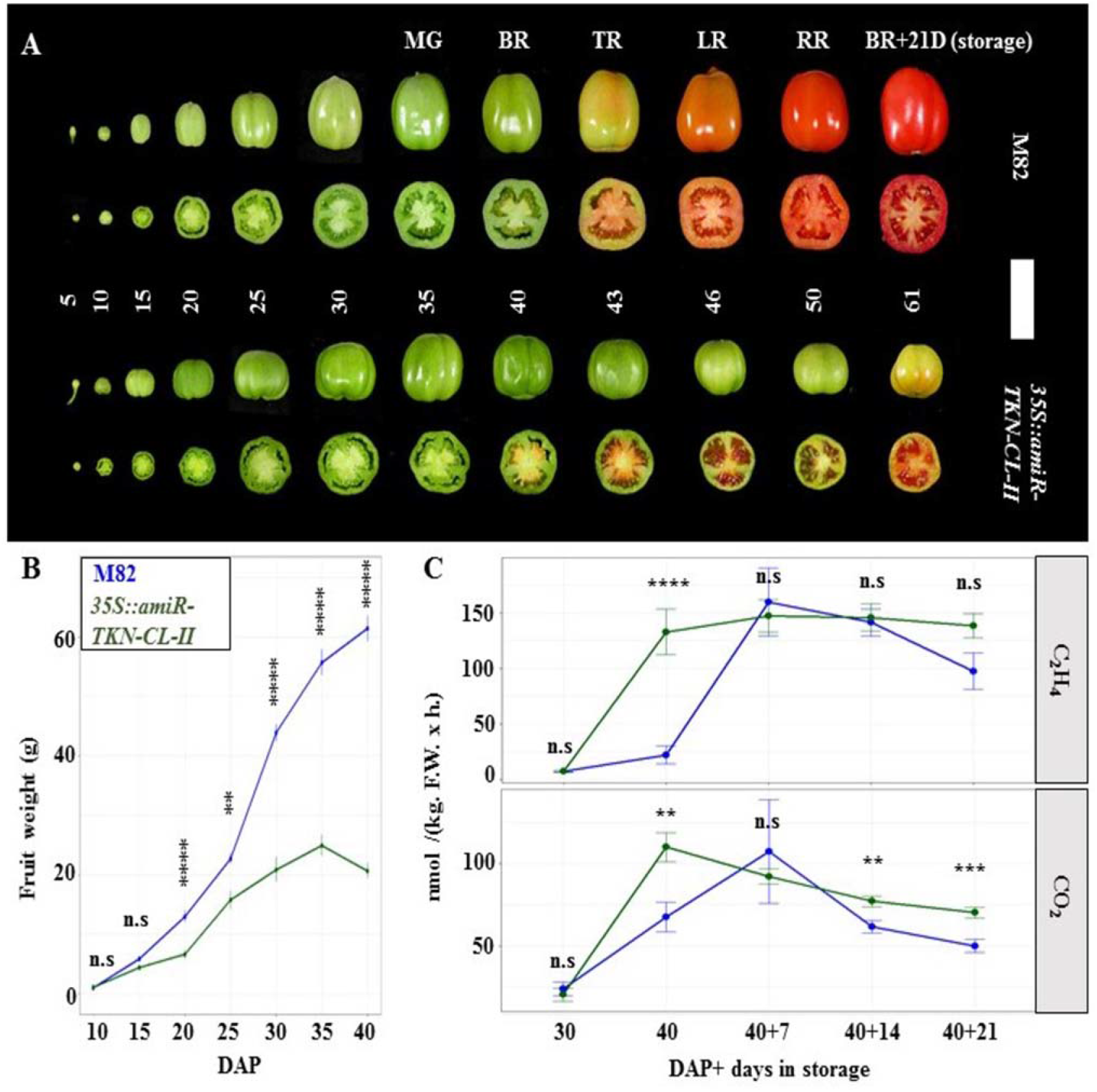
*Solanum lycopersicum CLASS-II KNOX* genes regulate fruit size and ripening patterns. **A:** Whole and cut fruits of M82 and *CLASS-II KNOX* knockdown plants at successive stages of development, demonstrating their different ripening patterns and fruit size; size-bar is 6cm; numbers indicate Days After Pollination (DAP); the letters on top indicate the ripening stages of the M82 fruits: **MG**-mature green; **BR**-Breaker, **TR**-Turning, **LR**-Light Red, **RR**-Red Ripen. Fruits harvested at 40 DAP (BR-stage) and stored for 21 days at shelf conditions are marked as **BR+21D. B:** Line plots displaying patterns of fruit weight (in gram (Y-axis)) accumulation in the developing (10 to 40 DAP (X-axis)) M82 (blue) and *CLASS-II KNOX* knockdown (*35S::amiR-TKN-CL-II*) (dark-green) fruits. Show significantly lower fruit weight accumulation in the transgenic fruits. **C:** Line plots displaying Ethylene (C_2_H_4_) (top panel) and Carbon dioxide (CO_2_) (bottom-panel) emissions (nano mol per kilogram of fruit weight an hour (Y-axis)) by the M82 (blue) and *CLASS-II KNOX* knockdown (dark-green) fruits at 30 and 40 DAP, and after 7,14 and 21 days of storage (of the 40DAP collected fruits).(40+7,40+14, 40+21) (X-axis). Show earlier C_2_H_4_ and CO_2_ emission peaks in the *CLASS-II KNOX* knockdown fruits. In **B-C:** Dots indicate the mean values; error bars indicate standard errors; Student’s t-test H_0_ probabilities are indicated as (n.s) (p-value)> 0.05; (*) (p-value) ≤ 0.05; (***) (p-value) ≤ 0.001; (****) (p-value) ≤ 0.0001; number of sampled fruits for each observation is ≥15.

Tomato is a climacteric fruit; thus, its transition to ripening is tightly associated with the elevated emission of ethylene and carbon dioxide (Fenn and Giovannoni 2020; Wang et al., 2020). To assess whether the unusual ripening patterns of the *SlCLASS-II KNOX* knockdown fruits are related to changes in their climacteric behavior, we measured the ethylene (C_2_H_4_) and carbon dioxide (CO_2_) levels emitted by the 30 and 40 DAP harvested fruits and by the 40 DAP harvested fruits that were then stored in controlled “shelf” conditions (22°C, 75% air-humidity) for 7,14 and 21 days. At 30DAP, low (close to 0) C_2_H_4_ levels were detected in both genotypes. At 40 DAP, approximately 25 and 130 fold elevation of ethylene levels were detected in the M7 82 and *35S::amiR-TKN-CL-II* fruits, respectively. After seven days of shelf–storage, the ethylene emission levels of the M82 fruits were increased to 150 nmol/(per kg(F.W.)x h) and became comparable to the levels measured in the 40 DAP *35S::amiR-TKN-CL-II* fruits. Both genotypes emitted similar C_2_H_4_ levels during the later storage stages (Fig. 2C – top panel).

The 30DAP fruits of both genotypes emitted low CO_2_ levels. At 40 DAP, the CO_2_ emission of both genotypes was elevated. However, the CO_2_ emission of the *35S::amiR-TKN-CL-II* fruits was significantly higher than the emissions of M82 fruits. M82 CO_2_ fruits emissions were elevated to the *35S::amiR-TKN-CL-II* levels after 7days of shelf-storage but were lower than in the *35S::amiR-TKN-CL-II* fruits again at the 14 and 21 shelf-storage stages (Fig. 2C –bottom panel). These results indicate that *SlCLASS-II KNOX* knockdown fruits reach the climacteric peak earlier than the control, promoting an earlier pigmentation change in their locular tissues and suggesting that the *SlCLASS-II KNOX* activity controls the pericarp ripening downstream to ethylene.

### *SlCLASS-II KNOX* expression at the *2A11* promoter domain is required for pericarp ripening

Characterization of the *35S::amiR-TKN-CL-II* line indicates that the ubiquitous *Sl CLASS-II KNOX* knockdown differentially regulates spatial patterns of pericarp ripening (Fig. 1D; 2A). Next, we wanted to identify the specific developmental time window during which *SlCLASS-II KNOX* activity is required to control fruit ripening. To address this goal, we used the OP/LhG4 trans-activation system (Moore et al., 1998) to silence *SlCLASS-II KNOX* genes in different stages of fruit development. Firstly, three fruit-specific *promoter::LhG4* lines: *pCRC:: LhG4* (Fernandez et al., 2009), *p2A11:: LhG4* (Gupta et al., 2021), and *pE8::LhG4* (see methods) were crossed to *OP::NLSmRFP* reporter line, to characterize their expression timing and patterns. The F1 carpels and fruits were collected at five-day intervals and imaged under the fluorescent stereoscope starting at anthesis. In *pCRC>>NLS-mRFP* plants, the mRFP signal was detected at the abaxial domain of initiating carpels, the abaxial domain of anthesis stage carpels, and external pericarp layers of the 10-20 DAP fruits (Fig. 3A-top row; S4-top row). Promoter *2A11* induced NLS-mRFP signal was detected in the pericarp, septum, and placental domains of the 10-50DAP (RR) fruits (Fig. 3A-middle row; S4 –middle row). The NLS-mRFP signal was first induced by the promoter *E8* at the locular tissues of the “Breaker” (40-DAP) stage fruits and was gradually expanded to all fruit domains by the RR (50DAP) stage (Fig. 3A-bottom row; S4 – bottom row).

**Figure 3:**
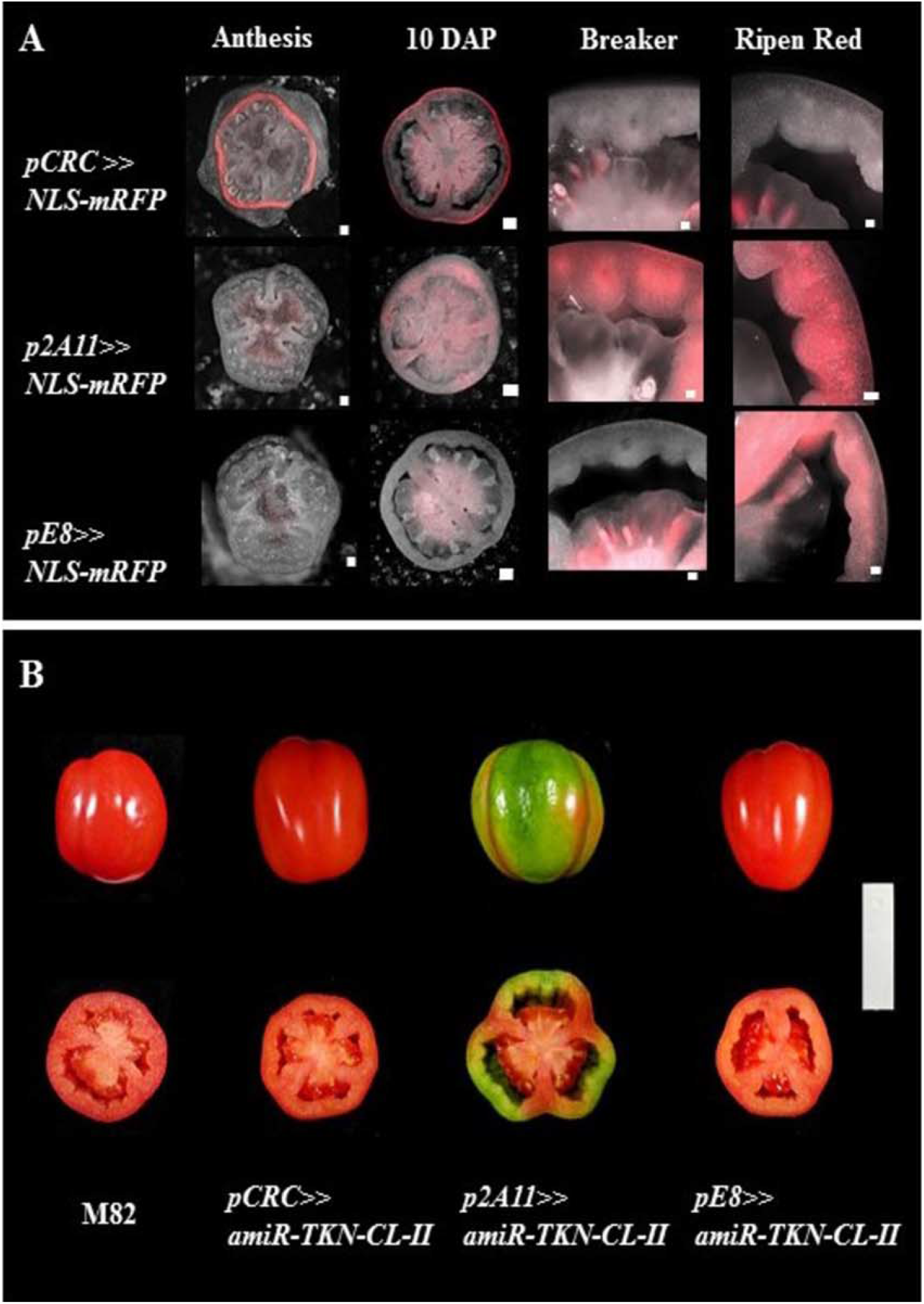
Stage-specific *CLASS-II KNOX* knockdown by 2A11 promoter delays pericarp ripening. **A:** Expression patterns of fruit-specific promoters used for developmental stage-specific knockdown of *CLASS-II KNOX* genes visualized by the *NLS-mRFP* signal (Red). Promoter CRC (Fernandez et al., 2009) top row, promoter 2A11(Gupta et al., 2021) middle row, and promoter E8 bottom row. The first column shows equatorial Anthesis-stage carpel sections (bar size is 100 µm), the second column shows equatorial 10 DAP fruit sections (bar size 500 µm), the third column shows sectors of equatorial “Breaker “ fruit sections (bar size is 500 µm), the fourth column shows sectors of equatorial “Ripen Red” fruit sections (bar size is 500 µm). **B:** Uncut 50 DAP fruits and equatorial fruit sections of M82 cultivar and trans-activation lines expressing the artificial micro RNA targeting CLASS-II KNOX genes under the control of the fruit-specific promoter characterized in **A**. The bar size is 6cm.

Trans-activation of the *OP::amiR-TKN-CL-II* construct by promoters *CRC* and *E8* did not result in any detectable fruit alterations. However, transactivation of the *OP::amiR-TKN-CL-II* with the *2A11 promoter* resulted in fruit ripening changes as observed with *35S::amiR-TKN-CL-II* fruits (Fig. 3B). This observation indicates that the pericarp ripening transition requires *Sl CLASS-II KNOX* activity at the domain covered by *promoter 2A11* temporal window.

### *RIPENING INHIBITOR* (*RIN*) activity promotes pigmentation change of the *Sl CLASS-II KNOX* knockdown locular tissues

The *ripening inhibitor* (*RIN*) gene, encoding a MADS-box protein, is a long investigated and well-established regulator of ripening (Klee and Giovannoni 2011; Li et al., 2018; Wang et al., 2020; Ito et al., 2017). The Ailsa Craig (AC) loss-of-function alleles were shown to delay the coloration of the internal fruit tissues (Ito et al., 2017; Ito et al., 2021). To examine the interaction of the *RIN* gene with *CLASS-II KNOX* genes in the regulation of tomato ripening, we used CRISPR-CAS9 to develop the M82 *cr-rin* knockout allele (Fig. S5). The M82 *cr-rin* allele has similar fruit phenotypes to previously described AC *CR*-*RIN* loss of function alleles (Ito et al., 2017; Ito et al.,2021); Its ripening progressed slower than the control, and even after 21 days of storage, the colors of the fruit locular tissues were yellow, whereas the pericarp coloration was similar to the control (Fig. 4A). The *cr-rin 35S::amiR-TKN-CL-II* knockdown fruits showed an additive effect; fruits with no pigmentation change in both the pericarp and locular tissues (Fig. 4A). In addition to pigmentation change, ripening is characterized by the softening of the pericarp tissues (Wang et al., 2018). We used a Stable Microsystem Texture Analyser system P2/N needle probe to measure the forces required for pericarp penetration. We found that the 40DAP (“Breaker”) stage *cr-rin* pericarp was slightly softer than the control, the *35S::amiR-TKN-CL-II* pericarp was slightly firmer, and the firmness of the *cr-rin 35S::amiR-TKN-CL-II* pericarps was not significantly different from M82 or from the *35S::amiR-TKN-CL-II* pericarp. After 21 days of shelf storage, the pericarp of M82 and *cr-rin* became 40-50% softer than at the beginning of storage, whereas the pericarp of *35S::amiR-TKN-CL-II* fruits have softened only by about 10-15% of its starting point. Strikingly, the fruits of *cr-rin 35S::amiR-TKN-CL-II* barely softened at all (Fig. 4B). These results suggest that *CLASS-II KNOX* genes and *RIN* have antagonistic functions during the initiation of ripening at the locular tissues, but both similarly promote ripening at the pericarp.

**Figure 4:**
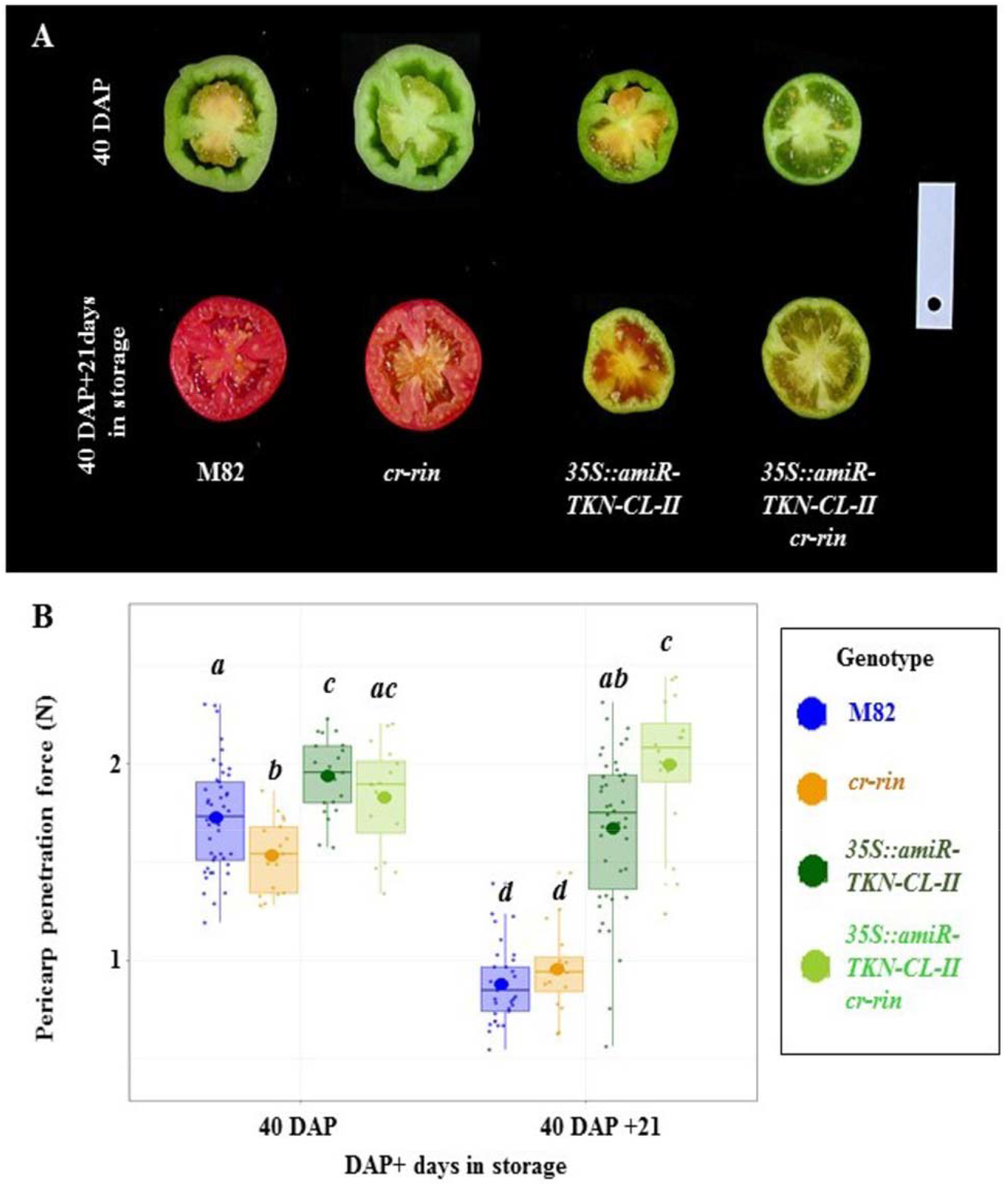
RIN activity is required for the ripening switch-induced pigmentation change of the *Sl CLASS-II KNOX* knockdown locular tissues. **A:** Images of the 40 DAP (top-row) and 40DAP collected then stored for 21 days in shelf conditions (22°C, 75% air-humidity) (bottom-line) M82, *cr-rin, 35S::amiR-TKN-CL-II* and *35S::amiR-TKN-CL-II/cr-rin* fruit equatorial sections, display yellow coloration of the locular domains in the *cr-rin* and *35S::amiR-TKN-CL-II/cr-rin* fruits and lack of pericarp pigmentation in the *35S::amiR-TKN-CL-II* and *35S::amiR-TKN-CL-II/cr-rin* fruits. The bar size is 6cm. **B:** Box plots displaying the distributions and average (large –dot) forces (in Newton (Y-axis)) required to penetrate the pericarps of the M82, *cr-rin, 35S::amiR-TKN-CL-II*, and *35S::amiR-TKN-CL-II/cr-rin* fruits at 40 DAP and after 21 days of shelf storage (X-axis). Different letters in italic indicate statistically significant (Student’s t-test p-value <0.05) means. Colors indicate different genotypes. The number of sampled fruits for each observation is ≥16.

## Discussion

### *Sl CLASS-II KNOX* genes regulate temporal and spatial patterns of fruit ripening

The initiation of ripening is considered to be coordinated with seed development and maturation (Klee and Giovannoni 2011; Wang et al., 2020). Indeed, the ripening of the fleshy fruits is usually initiated at the internal fruit domains, located in proximity to the seeds and gradually spreading into the pericarp tissues (Fig. **2A****-top panel**). Taxonomy studies of multiple Angiosperm families, including Solanaceae, describe numerous species with unusual flesh-fruit types, such as berries with hard pericarp (pepo or hesperidium), where the coloration and softening of the pericarp and internal fruit tissues are dissociated (Clausing et al., 2000; Knapp 2002). These observations indicate that in many species, fruit ripening and maturation are controlled both temporally and spatially. Although molecular mechanisms controlling fruit ripening have been thoroughly investigated (Klee and Giovannoni 2011; Li et al., 2018; Shinozaki et al., 2018; Fenn and Giovannoni 2020; Wang et al., 2020), little is known about its spatial regulation (Shinozaki et al., 2018).

Previous reports have demonstrated that *Arabidopsis thaliana CLASS-II KNOX* activity regulates the size of the silique and delays the initiation of senescence in dry fruits (Furumizu et al., 2015). Our results demonstrate that comparable fruit features are controlled by *CLASS-II KNOX* genes in the fleshy tomato fruits, where they regulate fruit size, delay the initiation of the climacteric peak and delay locular pigmentation (Fig. 2A-C). However, *SlCLASS-II KNOX* genes activity is also required to promote pericarp ripening (Fig. 2A;3B). Thus, *SlCLASS-II KNOX* genes have antagonistic functions in coordinating the ripening of the internal and external fruit domains (Fig. 5).

**Figure 5:**
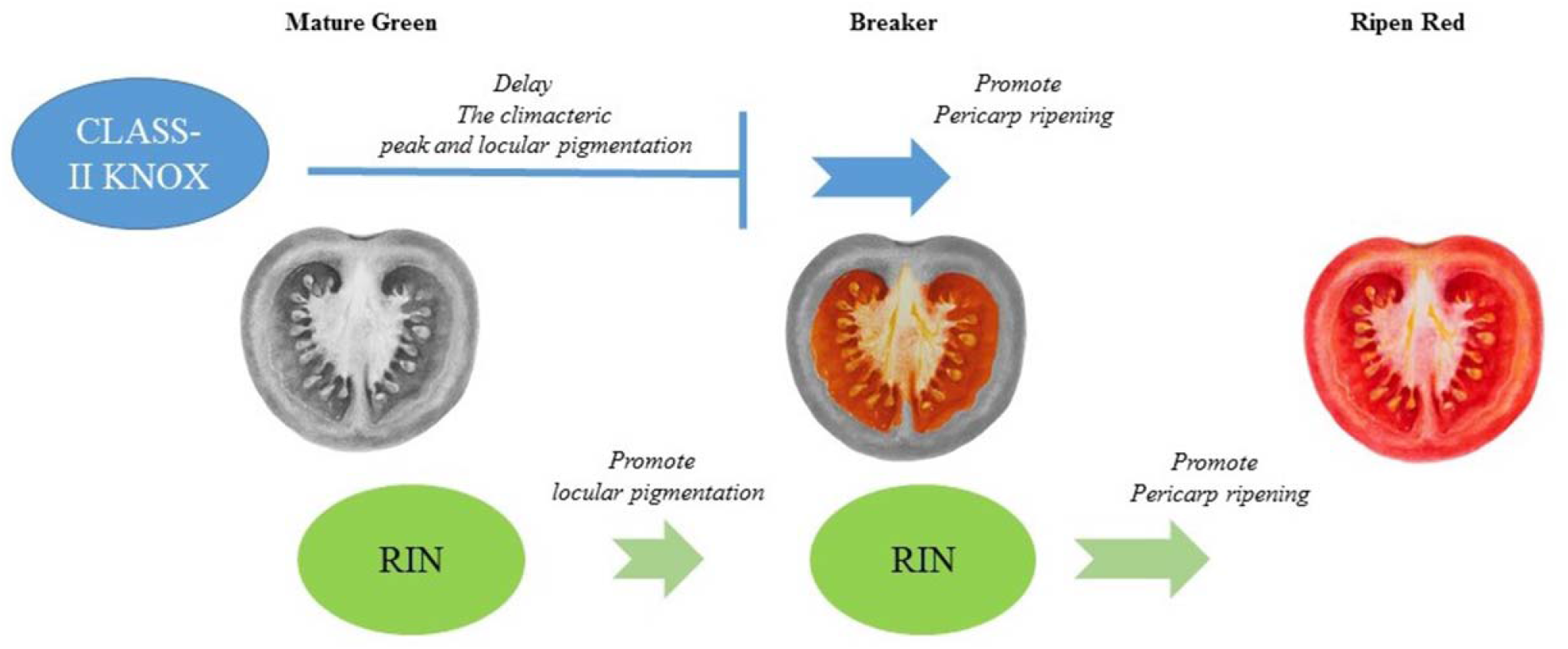
Proposed model for spatial control of tomato ripening by *CLASS-II KNOX* and *RIN* genes. Pre-breaker stage *CLASS-II KNOX* activity is required to delay the climacteric peak and pigmentation of locular tissues and to promote the pericarp ripening. *RIN* expression starts at the Mature Green stage and is required for locular pigmentation and promotion of pericarp ripening. The two genes have antagonistic functions during the initiation of ripening at the locular domain and similar functions in the coordination of the pericarp ripening. (Red colors indicate pigmentation of fruit domains).

### *SlCLASS-II KNOX* and *RIN* genes have antagonistic functions during ripening initiation at the locular domain and similar functions in the pericarp ripening

The role of the RIN gene in ripening has been rigorously studied (Klee and Giovannoni 2011; Li et al., 2018; Wang et al., 2020; Ito et al., 2017). Its expression is initiated at the Mature Green stage locular tissues and is tightly associated with ripening progression across all fruit domains (Shinozaki et al., 2018). Examination of the M82 *cr-rin* mutant fruits suggests that RIN activity promotes pigmentation switch of the locular fruit tissues (Fig. 4A). *35S::amiR-TKN-CL-II cr-rin* double mutants analysis suggests that the two gene systems act antagonistically during ripening of the locular tissues, when *CLASS-II KNOX* genes delay the climacteric peak and pigmentation of this domain whereas *RIN* promotes it (Fig. 2A;2C;4A;5). However, both gene systems are similarly involved in promoting the pericarp ripening (Fig. 4A-B; 5). These results suggest that molecular mechanisms regulating ripening are not uniform across different fruit domains.

Our observations advocate that further investigation and identification of the molecular networks regulated by the *CLASS-II KNOX* and *RIN* genes at the different fruit domains before and during ripening would be necessary steps for future studies aiming to pinpoint the genetic mechanisms governing ripening at different fruit tissues.

## Material and Methods

### Protein phylogenetic analysis

Sequences of CLASS-II KNOX proteins were aligned using muscle. Phylogenetic tree and bootstrap values were obtained using IQ-TREE v. 1.6.12 (parameters -alrt 1000 -bb 1000). The tree was visualized using iTOL (itol.embl.de). Branched with bootstrap values of less than 75/100 were collapsed.

### Plant material

M82 cultivar was used as a control and transformation background for the transgenic *35S::amiR-TKN-CL-II, promoter E8::LhG4, OP::amiR-TKN-CL-II* lines used in this study. The artificial micro-RNA constructs were generated as described in (Fig. S2) and by (Alvarez et al., 2006) using the 35S promoter (Fernandez et al., 2009) or OP/LhG4 system (Moore et al., 1998) in BJ36 plasmid (Eshed et al., 2001) same system was used to subclone 1095 bp sequence upstream of the *Solyc09g089580* (E8) to generate the *promoter E8::LhG4* driver. The transgenic tomato plants were generated using previously described protocols (McCormick 1997). The *OP::NLS::RFP, promoter CRC::LhG4, promoter2A11::LhG4* lines were previously described by (Shalit et al., 2009; Fernandez et al., 2009; Gupta et al., 2021), respectively.

### Development of the CRISPR-CAS9 mutant

The guide RNA sequences targeting the coding regions of *Solyc05g 012020* were designed using the benchling software (www.benchling.com). PCR primers with the guide sequences (Fig. S5) were synthesized by IDT(www.eu.idtdna.com) and used to amplify and modify constructs carrying the CRISPR-guide-RNA sequences (Brooks et al., 2014). The GOLDEN-GATE cloning system (Engler et al., 2014) was used for subcloning the gRNAs under the control of Arabidopsis U6 promoter (Dahan-Meir et al., 2018) into a binary plasmid containing SlUbiquitin10 promoter::CAS9 constructs (Dahan-Meir et al., 2018). After cloning, the plasmid was transformed into M82 plants, as described in the plant material section.

### Plant Growth conditions

Plants were grown at ARO Volcani temperature-controlled greenhouses during winter (September-April) and spring (March-June) seasons using the standard agronomic practice, under the average daily temperatures 25-30°C and average night temperatures 18-20°C. All described plants and phenotypes were grown, observed, and measured at least once during each season.

### Postharvest storage

Fruits were harvested at the “Breaker” 40DAP stage and stored in open carton boxes in the dark controlled climate chamber at 22°C, 75% air humidity. Once a week, the fruits were taken for ethylene and carbon dioxide measurements and weighted. After three weeks of storage, the fruits were weighed, assessed for ethylene, carbon dioxide, and pericarp stiffness (penetration force).

### Imaging

Expression of the NLS::RFP marker was assessed in the dissected flowers, ovaries, and fresh-cut fruits using Nicon SZM25 stereoscope; SHR Plan Apo 1X WD60 objective and 560nm excitation and 660 nm emission filters. The mature fruits were cut and imaged using a Canon Power Shot SX520 HS camera.

### Transcription levels assessment by the quantitive PCR test

All plant tissues used to assess the transcription levels were collected into liquid nitrogen 3-4 hours after dawn. The RNA was extracted using the Gene All, “RiboEx – total RNA isolation” kit (#301-001); the” DNA-free DNase Treatment &Removal” kit (Invitrogen ThermoFisher Scientific AM1906) was used to remove any DNA contaminations. Subsequently, one µg of RNA was used to prepare cDNA with the “High capacity RNA–to-cDNA” kit (#4387406 Applied biosystems-ThermoFisher Scientific). The qPCR analysis of the total cDNA was performed using the “Fast SYBR Green Master Mix “(# 4385612 Applied biosystems-ThermoFisher Scientific), gene-specific primers, and SAND (Expósito-Rodríguez et al., 2008 primers as control (Suppl. Tabel 1). The assay was performed with Applied Biosystems Step One Plus Real-time PCR system and software. The Relative expression values were calculated as described (Pfaffl 2004).

### Carbon dioxide and Ethylene measurements

Carbon dioxide was measured by a Gas Chromatograph (GC) equipped with a thermal conductivity detector TCD (Shimadzu Series 2014, Kyoto, Japan). The 1 m-long and 2.1 mm-internal diameter column was packed with Porapak N (Supelco, Bellefonte, PA, USA) and mesh size of 80/100.

Temperatures of the injector port, column, and detector were 90, 60, and 100°C, respectively. The concentrations were calibrated with a CO2 standard of 1.0 % (v/v). Helium was used as the carrier gas, and the flow rate was 20 mL min-1.

Ethylene concentration was determined with a GC equipped with a flame ionization detector FID (Varian 3300; Varian, Palo Alto, CA, USA) and a two m-long HayeSep T100/120 column (Grace Davison Discovery Sciences, Deerfield, IL, USA). The concentrations were calibrated with an ethylene standard of 1 ppm. The injector port, column, and detector temperatures were 75, 80, and 60°C, respectively. Helium was used as the carrier gas, and the flow rates were 30 mL min-1.

### Penetration force measurements

Stable Microsystem TA.XTplusC Texture Analyser, Godalming, UK (https://www.stablemicrosystems.com) with a P2/N needle probe was used to assess the pericarp penetration force at the “Breaker” stage and upon twenty-one days of storage.

### Calculations and statistical analysis

All plants were randomized in the net houses. Only the plants growing at the same season were compared to the controls growing at the same conditions. The means differences were compared by the Student’s –t-test, using the R-language, “tydiverse,” and “rstatix” packages.

## Accession Numbers

*SlKNAT3*: Solyc07g007120; *SlKNAT4:* Solyc12g010410; *SlKNAT5:* Solyc08g041820; *SlKNAT7:* Solyc08g080120; *RIN*: Solyc05g012020

## Supplemental Data

**Supplemental Figure S1:** Cube plots summary of the fruit domain and tissue-specific expression patterns of the Sl CLASS-II KNOX genes

**Supplementary Figure S2:** Artificial microRNAs targeting class II SlKNAT genes

**Supplementary Figure S3:** Bar plots indicating the leaf numbers produced by M82 and 35S::amiR-TKN-CL-II plants before the first flower.

**Supplementary Figure S4:** Detailed summary of the expression patterns of fruit-specific promoters used for developmental stage-specific knockdown of CLASS-II KNOX genes

**Supplementary Figure S5:** Gene model of *RIN Solyc05g012020* gene indicating the gRNAs and sites of the deletion mutation generated by the CRISPR-CAS9.

## Acknowledgments

We thank Yuval Eshed, Naomi Ori, and Ilan Paran for their valuable comments and manuscript discussion. We also thank Zlil Lazarovich for the visual art image used in Figure 5, Gil Yashuron, and Aharon Bellalou for the assistance at the greenhouse.

